# FlyAnatoView: Visualization of *Drosophila* Single-cell RNA-seq Datasets on Anatomographs

**DOI:** 10.64898/2026.09.28.755010

**Authors:** Yanhui Hu, Jenna Kalina, Austin Veal, Ben Ewen-Campen, Tarryn Porter, Julia Song, Yicheng Luo, Kerui Huang, Damien Goutte-Gattat, Nicholas H. Brown, Irene Papatheodorou, Norbert Perrimon

## Abstract

Single-cell RNA sequencing (scRNA-seq) is a powerful approach for characterizing gene expression at single-cell resolution and identifying distinct cell populations within complex tissues. However, tissue dissociation disrupts the spatial relationships among cells, making it difficult to determine their original anatomical locations and interpret gene expression patterns in the context of tissue organization. To address this limitation, we developed FlyAnatoView (https://www.flyrnai.org/tools/rna_seq_base/web/svg_files/), a resource that projects scRNA-seq data onto anatomical diagrams of *Drosophila melanogaster*. By integrating gene expression data with anatomical representations, FlyAnatoView helps restore spatial context and enables users to explore expression patterns across tissues, organs, and cell types. This resource provides an intuitive way to connect transcriptomic profiles with anatomical structures, facilitating the interpretation and exploration of scRNA-seq datasets.

## INTRODUCTION

Single-cell RNA sequencing (scRNA-seq) has become an essential tool in biological research, offering far greater resolution into the transcriptional landscape of individual cells than was previously possible. Unlike bulk RNA sequencing, which averages gene expression across an entire population of cells and thereby obscures important biological variation, scRNA-seq resolves cellular heterogeneity at single-cell level, revealing subpopulations, rare cell types, and transitional states that would otherwise go undetected. This technology has driven large-scale community efforts such as the Human Cell Atlas (Regev *et al*. 2017), which catalogs cell types across tissues and organisms, and the Fly Cell Atlas (FCA), which aims to comprehensively catalog cell types across the entire adult fly (Li *et al*. 2022). Note that in this study, we do not distinguish between single-cell RNA sequencing (scRNA-seq) and single-nucleus RNA sequencing (snRNA-seq); for simplicity, we use the term scRNA-seq throughout to refer to both.

scRNA-seq profiles the gene expression of individual cells, but it requires dissociating tissue into a single-cell suspension, a process that destroys all information about where those cells originally resided. Spatial transcriptomics addresses this limitation by mapping RNA transcripts directly within intact tissue sections, adding the physical and structural context that scRNA-seq alone cannot provide (Piwecka *et al*. 2023; Jain and Eadon 2024). This context is especially important because many tissues exhibit continuous or graded gene expression across space, such as along the segments of the kidney nephron, and such gradients are invisible in a purely transcriptional embedding (e.g., UMAP or t-SNE).

Although spatial transcriptomics is widely used in vertebrates, it has been applied far less often in *Drosophila*, and the tools available for restoring spatial context to expression data in flies remain limited. Existing resources are restricted either by developmental stage or by tissue. FlyExpress (Kumar et al. 2011; Kumar et al. 2017) and the Berkeley Drosophila Genome Project (BDGP) (Tomancak et al. 2002; Tomancak et al. 2007) provide spatial context for gene expression through curated in situ hybridization images, but they predate the scRNA-seq era and cover only embryogenesis. Virtual Fly Brain (VFB) (Milyaev et al. 2012) is a richly curated hub for *Drosophila* neuroanatomy that integrates 3D image stacks, neuron morphologies, connectomic data, and gene/transgene expression patterns onto standardized brain templates, and it includes tools for browsing single-cell transcriptomic data registered to these templates. However, its scope is limited to the central nervous system.

In other species, efforts to restore spatial context to expression data are more developed but still have constraints. The Expression Atlas “anatomogram” tool (Papatheodorou et al. 2018) projects bulk and single-cell expression data onto adult human and mouse body diagrams, allowing users to visually query where a gene is expressed across organs and tissues. Its coverage is limited to human and mouse, and its resolution is generally at the level of whole organs rather than individual cell types within an organ’s substructure. Building on these anatomograms, gganatogram (Maag 2018) repurposes the same underlying Scalable Vector Graphics (SVG) diagrams to enable programmatic, reproducible visualization of user-supplied expression data through the ggplot2 framework. Although it has been extended to 28 organisms, its species coverage does not include *Drosophila*.

Together, these resources highlight a clear need to extend the anatomogram concept to adult *Drosophila* organ systems using modern single-cell/single-nucleus transcriptomic datasets. To address this need, we developed FlyAnatoView, a resource for visualizing scRNA-seq-derived gene expression directly on anatomically accurate diagrams. This resource covers portals for the major organs of the adult fly, including the dorsal vessel, digestive system, Malpighian Tubules (MT), male and female reproductive systems, and trachea, as well as a whole-body portal. The FCA dataset (Li *et al*. 2022) was used for the dorsal vessel, digestive system, and male reproductive portals, while datasets from other published studies (Rust *et al*. 2020; Xu *et al*. 2022; Li *et al*. 2024; Xu *et al*. 2024; Liu *et al*. 2025; Ewen-Campen *et al*. 2026) were used for the MT, female reproductive, tracheal, and whole-body portals. Users can query the expression of a single gene or compare two genes simultaneously on any portal; a subset of the portals additionally allow comparison of expression across different genotypes.

## MATERIALS AND METHODS

### Generation of anatomograms and identification of the corresponding datasets

Anatomical illustrations of various *Drosophila* organs including the dorsal vessel, midgut, renal system (Malpighian Tubules), and male and female reproductive systems, were created in SVG format using Inkscape (https://inkscape.org/), based on reference illustrations from textbooks and the primary literature. Sections and cell types were annotated according to the FlyBase anatomy ontology. The tracheal, proventriculus and whole-body illustrations, by contrast, were designed and annotated directly based on the cell types identified in their corresponding scRNA-seq datasets.

### Reshaping scRNA-seq datasets for anatomograms

The Fly Cell Atlas (FCA) dataset (Li *et al*. 2022) was downloaded as a loom file from https://www.flycellatlas.org/#data, and its 250 annotated cell types were manually grouped to broad terms spanning 14 biological systems. Cell types belonging to organs covered by our anatomograms were selected and manually mapped to the corresponding anatomical ontology terms; for example, digestive-system cell types in the FCA dataset were matched to the anatomical illustration of the midgut. Because the resolution of cell-type annotation in a given dataset does not always match that of the corresponding illustration, mappings could be one-to-one, one-to-many, or many-to-one.

Additional datasets for the renal system, female reproductive system, trachea, and whole body were obtained from the Gene Expression Omnibus (GEO) repository (Table 1). Cell types annotated in each dataset were manually matched to the ontology terms used in the corresponding illustration. Where multiple candidate datasets existed for a given organ, the dataset with the greatest number of successfully matched cell types was selected for use in that portal.

**Table 1.**
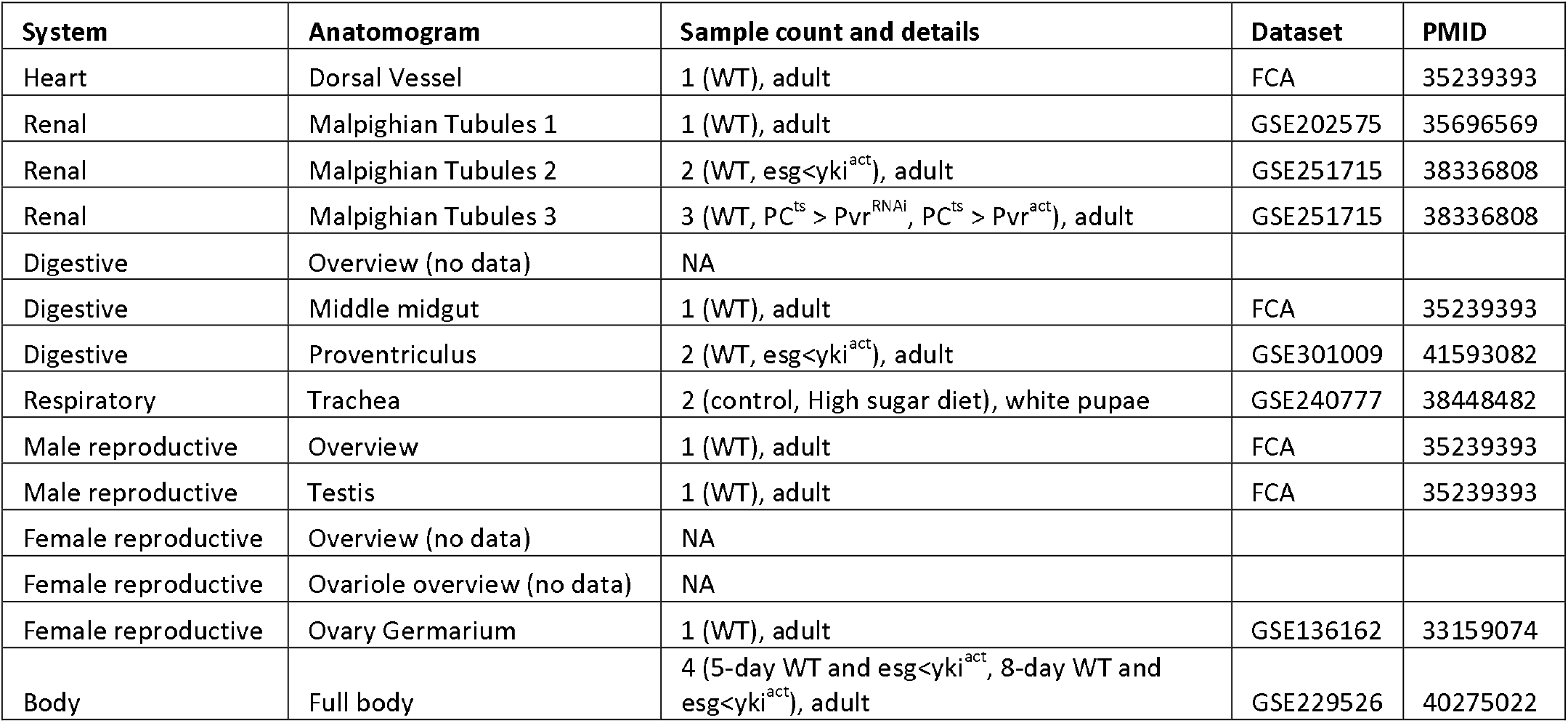
FlyAnatoView coverage and data source.

For each cell type represented in an illustration, raw UMI counts were summed across all cells of that type to generate a single pseudo-bulk count vector, which was then normalized to counts per million (CPM) to account for differences in cell number and sequencing depth across samples; average counts and percent-expression values were calculated in parallel. This pseudo-bulking strategy naturally accommodates one-to-many mappings, in which a single cell type on the anatomical illustration corresponds to multiple cell types in the dataset, by aggregating counts across all corresponding dataset cell types. For many-to-one mappings, in which a single dataset cell type corresponds to multiple regions on the illustration, the same expression values are applied to each of the corresponding illustrated cell types. Code for this data-reshaping pipeline is available on GitHub (https://github.com/jennakalina/anatomograms).

### Development of the online resource

The FlyAnatoView web application was developed in PHP using the Silex micro-framework, built on Symfony components, with JSON used to exchange data between client and server. The front end uses HTML for document structure, JavaScript and jQuery for client-side interactivity, and Twig as a templating engine for generating pages from application data. Bootstrap supports interface styling and responsive design, and DataTables enhances the presentation and interactivity of tabular results. The resource is publicly accessible at www.flyrnai.org/tools/rna_seq_base/web/svg_files. Gene symbol and gene identifiers entered by user goes through the GeneID mapping tool developed for FlyRNAi (Hu *et al*. 2017; Hu *et al*. 2021; Hu *et al*. 2025).

## RESULTS AND DISCUSSION

*Drosophila* has long served as a well-established model organism in biological research (Ozturk-Colak *et al*. 2024), and anatomical illustrations of its various organs are available from numerous sources, including textbooks and primary research articles. The ovary, for example, is a particularly well-characterized organ, with illustrations available at multiple levels of resolution down to individual cell types (King 1970). Building on these existing resources, we generated anatomical illustrations of the *Drosophila* ovary in SVG format, drawing on textbook and literature references, and annotated the resulting images with identifiers from the FlyBase anatomy ontology (Figure 1). We applied the same approach to generate illustrations of the following adult organs: midgut, male reproductive system, dorsal vessel, and renal system (Malpighian Tubules). In contrast, illustrations of the tracheal system, the proventriculus and whole-body illustrations were designed and annotated directly from the cell types identified in their corresponding single-cell datasets (Li *et al*. 2024; Liu *et al*. 2025; Ewen-Campen *et al*. 2026).

**Figure 1.**
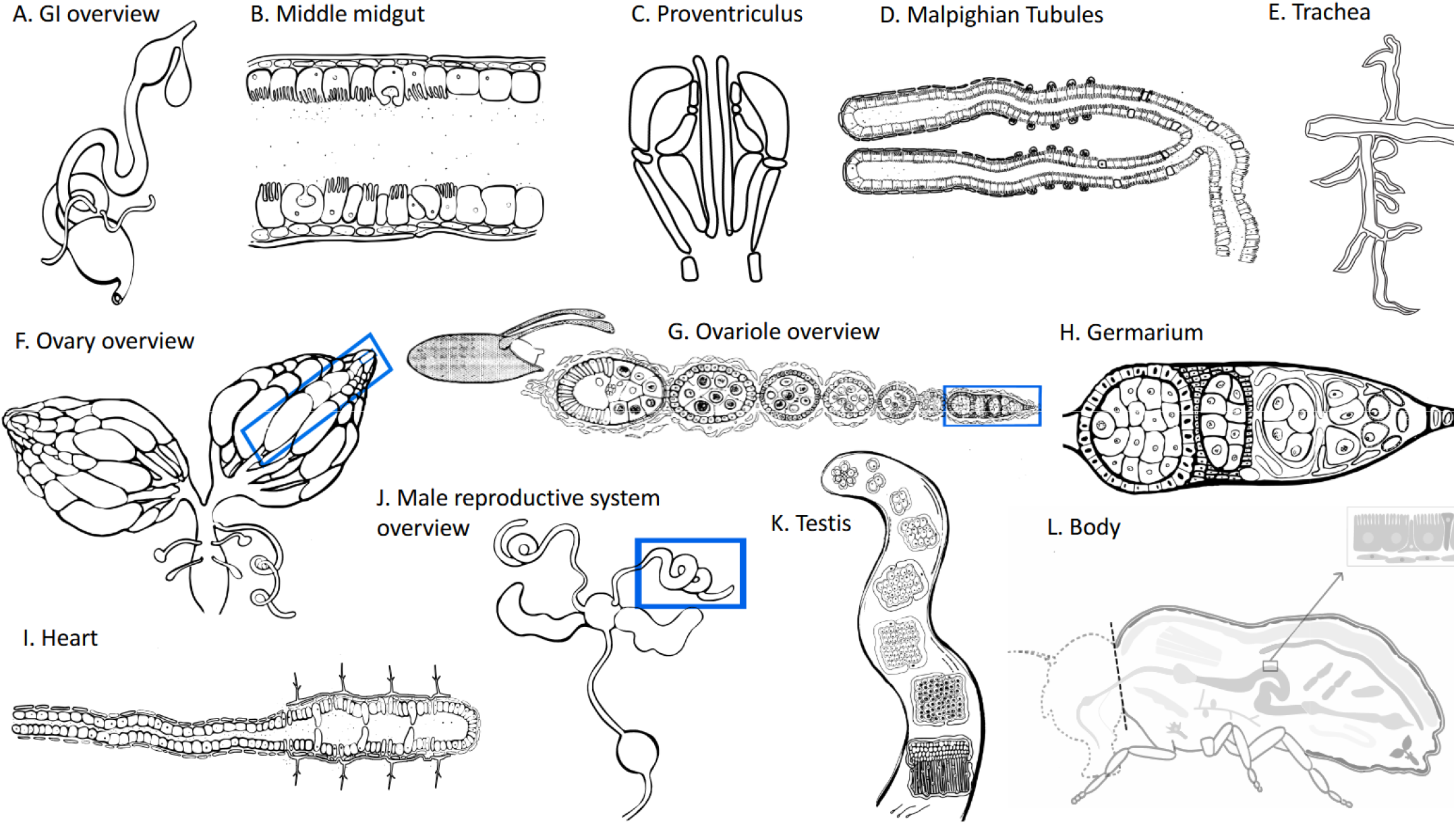
Anatomograms. (A) Digestive System Overview, (B) Middle midgut, (C) Proventriculus, (D) Malpighian Tubules, (E) Trachea, (F) Female reproductive system (the big blue box indicates one single Ovariole while the small blue box indicates one single Germarium), (G) Ovariole (the blue area indicates Germarium), (H) Germarium, (I) Dorsal Vessel, (J) Male reproductive system (the blue area indicates Testis), (K) Testis, (L) Body.

Various scRNA-seq datasets (Rust *et al*. 2020; Li *et al*. 2022; Xu *et al*. 2022; Xu *et al*. 2024), including the FCA dataset, were processed and reshaped to best match the resolution of the anatomical illustrations. Where multiple candidate datasets existed for a given organ system, the dataset with the greatest number of cell types successfully matched to the illustration was selected (Table 1).

We developed an online resource comprising multiple portals for mining these curated scRNA-seq datasets (Figure 2). The landing page presents a menu of all available illustrations, organized by organ systems (Figure 2A). For each portal, users can search for a single gene or compare two genes side by side (Figure 2B), and can select percent expression, average count, or CPM from a dropdown menu as the expression parameter of interest. Expression values are color-coded on the anatomogram, with color intensity reflecting the expression level for the selected parameter; users can also customize the display colors (Figure 2C). A table summarizing expression statistics is displayed alongside each illustration, along with links to the original publication and the corresponding UMAP/t-SNE-based data portal (Figure 2C). In addition, four of the portals allow users to compare gene expression across genotypes (Table 1). For example, one of the Malpighian Tubule portals allows users to compare expression between wild-type flies and flies bearing tumors in their gut induced by the expression of an activated form of Yorkie in the intestinal stem cells (ISCs), referred to as “gut yki tumors” (Figure 3). Users can download the anatomogram images rendered with the expression values of the queried gene, along with their accompanying legends in SVG format.

**Figure 2.**
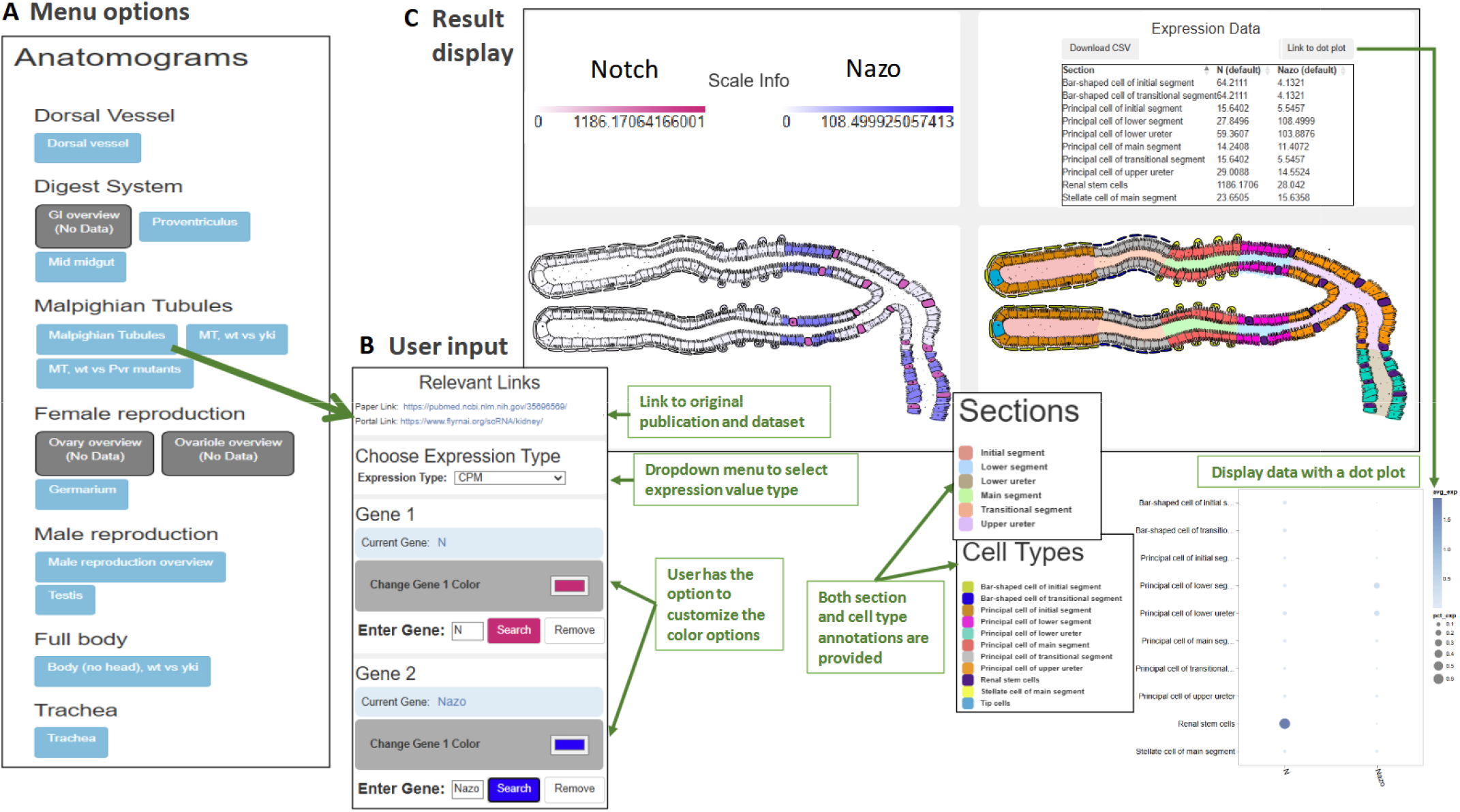
User interface of FlyAnatoView, illustrated using the Malpighian Tubules portal. (A) The landing-page menu lists anatomograms organized by organ system (Dorsal Vessel, Digestive System, Malpighian Tubules, Female/Male reproductive system, Full body, Trachea); grayed-out buttons marked “(No Data)” denote overview illustrations with no associated expression data. (B) The user-input panel, reached by selecting a portal (e.g., “Malpighian Tubules”), provides links to the original publication and data portal, a dropdown to select the expression metric (CPM, average count, or percent expression), and search fields for one or two genes (here, *Notch* and *Nazo*), each with a customizable display color. (C) The result display shows the two selected genes’ expression levels overlaid on the Malpighian Tubule anatomogram, color-coded by intensity (scale bars indicate the expression range for each gene), alongside a sortable data table listing expression values for every annotated section and cell type. Section and cell-type identities corresponding to the colored regions on the anatomogram are provided in the accompanying legends. Above the data table, a link is provided for user to visualize the data with a dot pot.

**Figure 3.**
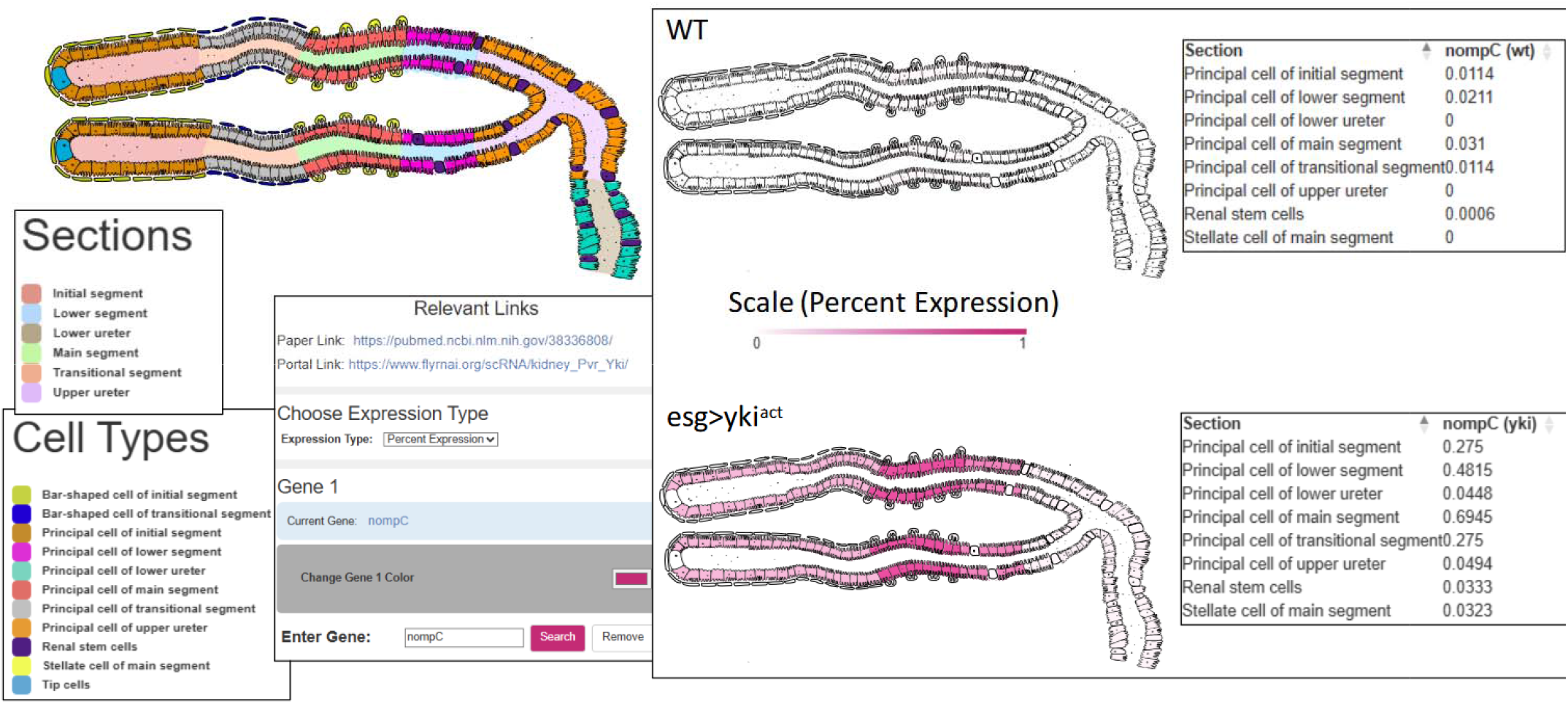
Cross-genotype comparison. Expression of *nompC* (*no mechanoreceptor potential C*) is displayed on the Malpighian Tubules anatomogram side-by-side for wild-type vs. yki-tumor samples, using “Percent Expression” as the metric. The data tables show expression values for each section (e.g., principal cells of the initial, lower, main, and transitional segments) with values visibly higher across nearly all sections in the yki sample compared to wild-type (e.g., principal cell of the main segment: 0.031 in WT vs. 0.6945 in yki), illustrating how the tool can visually and quantitatively reveal tumor-associated expression shifts region-by-region.

We demonstrated a representative use case on the midgut anatomogram portal by comparing the expression of *escargot* (*esg*) and *Allatostatin A* (AstA), two well-established marker genes that label distinct cell populations within the adult *Drosophila* midgut (Figure 4). *esg* is preferentially expressed in ISCs and their immediate daughter cells, enteroblasts (EBs), marking the progenitor compartment responsible for continual renewal of the midgut epithelium (Micchelli and Perrimon 2006; Ohlstein and Spradling 2006; Hung *et al*. 2020). In contrast, *AstA* is preferentially expressed in a subset of enteroendocrine cells (EEs), a rare, terminally differentiated population that secretes peptide hormones regulating gut motility and feeding behavior (Veenstra *et al*. 2008; Guo *et al*. 2019; Hung *et al*. 2020).

**Figure 4.**
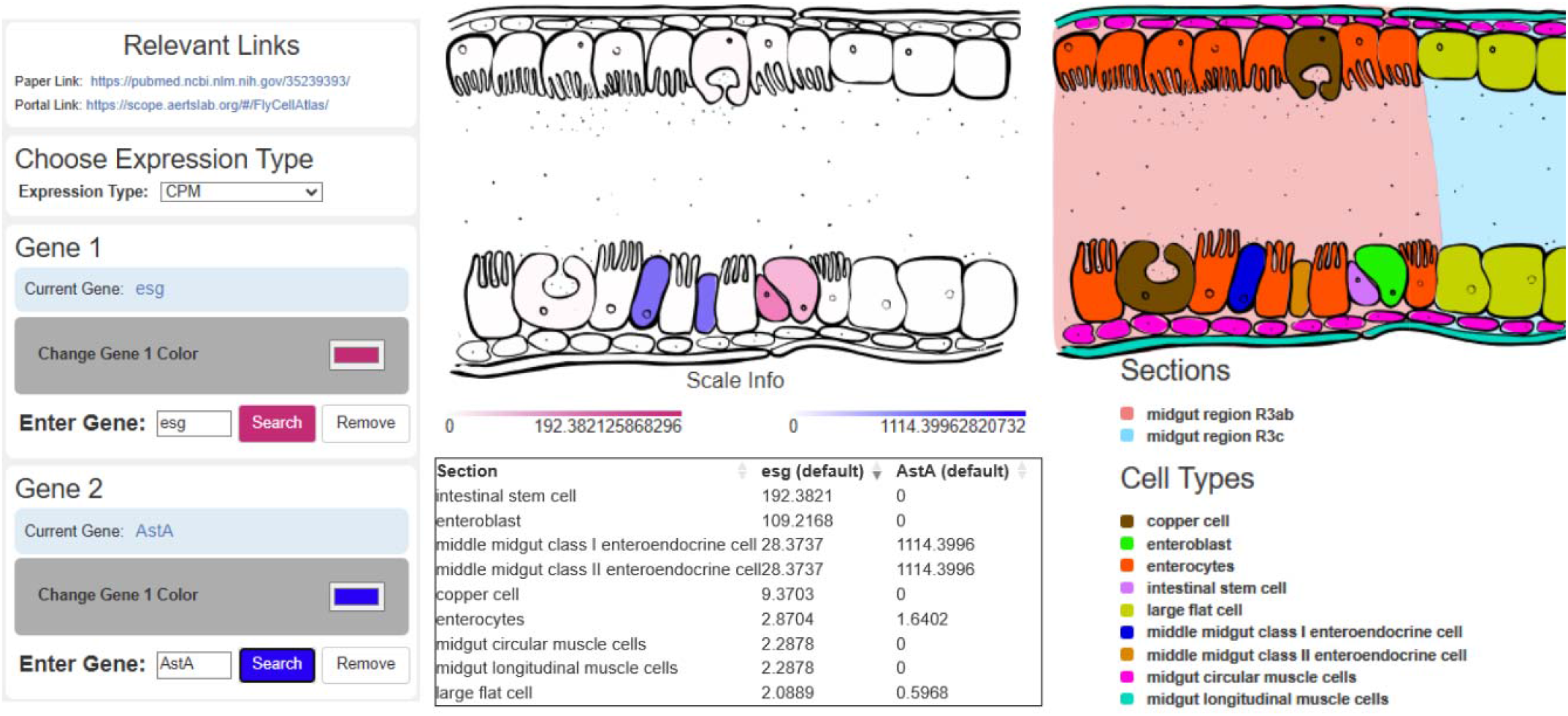
A representative use case of the midgut anatomogram: comparing the expression of *escargot (esg)* and *Allatostatin A (AstA)*. *esg* is preferentially expressed in intestinal stem cells (ISC) and enteroblasts (EB), whereas *AstA* is preferentially expressed in enteroendocrine cells (EE). The anatomogram clearly visualizes this cell-type-specific expression in its native spatial context.

Because ISCs/EBs and EEs occupy distinct positions and abundances along the midgut epithelium, this gene pair provides a clear test of whether cell-type-specific expression patterns are faithfully preserved when mapped back onto anatomical space.

When visualized side-by-side on the midgut anatomogram, *esg* and *AstA* showed clearly distinguishable, non-overlapping spatial signatures corresponding to their respective cell types, with each gene’s expression restricted to the anatomically annotated regions representing ISC/EB and EE populations, respectively. This result confirms that the pseudo-bulking and cell-type-to-anatomy mapping strategy underlying FlyAnatoView correctly preserves biologically meaningful expression differences, and illustrates how the tool allows users to directly and intuitively visualize where in the tissue a gene of interest is expressed—information that is not readily apparent from a UMAP or t-SNE embedding alone. anatomogram illustration clearly visualized the cell-type preference in its native spatial context.

In summary, FlyAnatoView provides a simple, accessible way to visualize *Drosophila* scRNA-seq gene expression data in its native spatial context, restoring the anatomical information lost during single-cell dissociation. By combining curated anatomical illustrations with a pseudo-bulking strategy that reshapes existing datasets to match illustration resolution, the resource allows users to query and compare gene expression across tissue regions, cell types, and even genotypes for select organs, all through an intuitive web interface. Currently spanning ten major organ systems and eight underlying datasets, FlyAnatoView complements existing single-cell atlases such as the Fly Cell Atlas by offering a spatially intuitive alternative to UMAP- or t-SNE-based views. We anticipate that this resource along with UMAP/t-SNE based resources will be useful to the *Drosophila* research community for generating hypotheses about tissue-specific gene function.

## ACKNOWLEDGEMENTS

We would like to thank the members of Perrimon laboratory, FlyBase consortium and EMBL-EBI for helpful suggestions regarding the project. We extend our gratitude to the Harvard Medical School Research Computing and IT-Client Services teams for their consultation, web hosting, and support. This work was supported by a joint NSF/BBSRC-BIO grant between Harvard Medical School, Cambridge University and the European Bioinformatics Institute (DBI-2035515, BB/T014563/2 and BB/T014008/1). N.P. is an investigator of Howard Hughes Medical Institute.

## DATA AVAILABILITY STATEMENT

The resource is publicly accessible at www.flyrnai.org/tools/rna_seq_base/web/svg_files. Code for this data-reshaping pipeline is available on GitHub (https://github.com/jennakalina/anatomograms).

